# Active Receptor Tyrosine Kinases, but not Brachyury, are sufficient to trigger chordoma in zebrafish

**DOI:** 10.1101/482687

**Authors:** Gianluca D‘Agati, Elena María Cabello, Karl Frontzek, Elisabeth J. Rushing, Robin Klemm, Mark D. Robinson, Richard M. White, Christian Mosimann, Alexa Burger

## Abstract

The aberrant activation of developmental processes triggers diverse cancer types. Chordoma is a rare, aggressive tumor arising from transformed notochord remnants. Several potentially oncogenic factors, including several receptor tyrosine kinase (RTK) genes, have been found deregulated in chordoma, yet causation remains uncertain. In particular, sustained expression of the developmental notochord transcription factor Brachyury is hypothesized as key driver of chordoma; nonetheless, experimental evidence for an oncogenic role of Brachyury in chordoma and its prognostic impact remains missing. Here, we developed and applied a zebrafish chordoma model to identify the notochord-transforming potential of implicated genes *in vivo*. We find that Brachyury, including a form with augmented transcriptional activity, is insufficient to initiate notochord hyperplasia. In contrast, the chordoma-implicated RTKs EGFR and KDR/VEGFR2 are sufficient to transform notochord cells within two to five days of development. Transcriptome and structural analysis of transformed notochords revealed that the aberrant activation of RTK/Ras signaling attenuates processes required for notochord differentiation, including of the unfolded protein response. Our results provide first *in vivo* indication against a tumor-initiating potential of Brachyury in the notochord, and imply activated RTK signaling as possibly initiating event in chordoma. These results provide a mechanistic framework for the pursuit of chemotherapeutic compounds to combat this aggressive tumor type.

## Introduction

Different genetic lesions can result in the malignant transformation of individual cells, ultimately leading to cancer. In particular, the reactivation or overexpression of developmental transcription factor genes has been repeatedly uncovered in a variety of tumors. Constitutive activation of developmental signaling pathways is even more common, in particular by activating mutations and amplifications of receptor tyrosine kinase (RTK) genes including *EGFR, FGFR, or VEGFR*. The permissive and instructive functions of these factors deployed in the embryo, reactivated or maintained unchecked in adult tissues, endow transformed cells with malignant properties of hyperproliferation, stemness, and tissue invasion. Nonetheless, the relationship between developmental regulators and maintenance of lineage identity for a given cancer’s cell of origin remains challenging to decipher.

Chordoma is a rare, slow-growing tumor typically occurring at the base of the skull or in sacrococcygeal regions along the spine (OMIM #215400). Chordomas are thought to result from malignant transformations of remnant cells of the embryonic notochord, a cartilaginous structure of mesodermal origin that supports embryo axis formation and spinal column formation before regressing during development (*1*). Despite its proposed origin from embryonic notochord cells, chordoma is typically first diagnosed in adults between ages 40 to 70, with an annual incidence of one case per million per year. Surgical removal of the tumor remains the most effective therapeutic option. Nonetheless, due to the deep tissue localization of chordomas, surgery of the strongly radio- and chemoresistant tumor is highly delicate or, in individual cases, even impossible. In addition, first treatment frequently results in re-occurring local tumors and possible metastases even after seemingly successful removal of the initial lesion. No proven systemic therapies are available for patients with re-occurring or non-resectable disease, nor against the ultimately fatal distant metastases (*2*).

To date, no single molecular pathway can be assigned as a fundamental chordoma-driving mechanism. Genomic sequencing of patient chordoma samples continues to uncover mutated genomic loci including *PTEN* (*3*), the Tuberous Sclerosis Complex (*TSC*) genes (*4*), *TP53* (*5*), and *BCL6* (*6*). Further, chordoma samples repeatedly feature increased expression, amplifications, or possibly activating mutations in several receptor tyrosine kinase (RTK) genes, including *EGFR* and the chromosome *4q12* genes *KDR/VEGFR2, KIT*, and *PDGFRA2* that relay their activity through Ras (*7*–*10*). Emphasizing the potential significance of reoccurring RTK involvement, EGF pathway inhibitors have been shown to curb the growth of chordoma cell lines and xenografts (*11*). Clinical trials have been performed using a variety of RTK inhibitors, including imatinib, sunitinib, and *EGFR* inhibitors, as well as the mTOR inhibitor rapamycin combined with imatinib (*2*).

A seemingly consistent feature of chordoma is the expression of the T-box transcription factor *T/Brachyury* (*12*). During development, *Brachyury* influences the formation of individual mesodermal fates, and is in particular an evolutionarily conserved regulator of the notochord (*13*). Intriguingly, the genomic locus encoding for *Brachyury* has been found to be duplicated or further amplified in familial and sporadic chordoma cases (*12, 14*). This data, together with detectable *Brachyury* expression in the vast majority of chordomas, points to a tumor origin from transformed remnant notochord cells. The transient nature of the notochord, which disappears before birth, and persistence of notochord cells in chordoma suggest that malignant transformation could depend on an early developmental pathway that maintains a notochord or early mesodermal program in remnant notochord cells. *Brachyury* expression is also increasingly reported from other cancers of the lung, small intestine, stomach, kidney, bladder, uterus, ovary, and testis, and its expression correlates with epithelial-to-mesenchymal transition (EMT), maintenance of an undifferentiated state, resistance of lung cancer cells to EGFR inhibition (*15*). *Brachyury* is suggested to inhibit the cell cycle by downregulating *CCND1 (*CyclinD), *RB (*pRb), and *CDKN1A (*p21), ultimately decreasing the susceptibility of tumor cells to cytotoxic therapies (*16*). Nonetheless, recent notochord-specific knockdown of *Brachyury* in mouse revealed that its activity is dispensable for notochord proliferation and EMT, questioning the potency of *Brachyury* mis- or overexpression alone to mediate chordoma formation (*17*). Consequently, the influence of *Brachyury* expression on chordoma initiation and tumor maintenance *in vivo* remains speculative, as are the pathways that maintain the notochord in an oncogenic progenitor state.

The lack of a unifying mechanism leading to chordoma formation necessitates animal models that recapitulate key aspects of the disease. Given the developmental origins of chordoma, the zebrafish offers a unique opportunity to dissect the mechanisms of tumor initiation and progression. Controlled by regulators including Brachyury that drive deeply conserved notochord-forming mechanisms across vertebrates, the early notochord is principally formed by central vacuolated cells that provide hydraulic stability. In zebrafish, Notch-signaling dependent differentiation results in an epithelial layer of outer sheath cells (*18*). The sheath cells surround the vacuole cells and secrete an extracellular matrix (ECM) composed of collagens, laminins, and proteoglycans that encapsulate the notochord. Work in medaka has established that endoplasmic reticulum (ER) stress occurs physiologically during this process, which requires the unfolded protein response (UPR) transducers Atf6 and Creb3l2 for the proper alignment of the notochord cells and for export of type II collagen (*19, 20*). Notochord formation in Xenopus has also been linked to the progressive activation of UPR via Xbp1 and Creb3l2 to drive its differentiation (*21*). Subsequently, the notochord ossifies to form the spine segments, while remnant notochord cells turn into the gel-like nucleus pulposus inside the intervertebral disks of the spinal column (*22, 23*).

We recently established in zebrafish the first animal proxy for chordoma onset based on notochord-specific expression of *HRASV12* that recapitulates oncogenic RTK/Ras pathway activation (*24*). Our model uses the bimodal *Gal4/UAS* system, in which a notochord-specific transgene expresses the *Gal4* transcription factor that drives a separate candidate transgene with *Upstream Activating Sites (UAS)*. With virtually 100% animals affected as early as 2-3 days post-fertilization (dpf), *HRASV12*-expressing embryos rapidly develop prominent notochord hyperplasia that shares key histological features with human chordoma samples (*24*).

Here, we applied this *in vivo* platform to test the potency of chordoma-implicated factors in driving notochord hyperplasia. We established transgenic zebrafish for *col2a1aR2:KalTA4* for robust injection-based candidate gene expression with fluorescent tags in the developing notochord. *col2a1aR2*-driven transient mosaic expression of *HRASV12* triggered chordoma-like notochord hyperplasia akin to stable genetic insertions. Notably, notochord-specific overexpression of human or zebrafish *Brachyury*, including a version with augmented transcriptional activity, failed to cause notochord hyperplasia in the observed timeframe of 5 dpf. In contrast, overexpression of the chordoma-implicated RTKs *EGFR* and *KDR/VEGFR2* potently induced a chordoma phenotype. Transcriptome sequencing and cellular ultrastructure analysis uncovered that Ras-mediated RTK signaling drives excessive secretory pathway activity, extracellular matrix (ECM) buildup, and a suppression of the Unfolded Protein Response (UPR) in transformed notochord sheath cells. These processes hinder the transformed sheath cells from further differentiation, yet ultimately trap the hyperproliferating mass with its accumulating ECM. Taken together, our data indicate that *Brachyury per se* might be insufficient to initiate chordoma and rather reflects the maintained notochord lineage identity of the transformed cells. Our results instead propose that aberrant RTK signaling, possible by activation of several repeatedly chordoma-implicated RTK genes (*7*–*10*), presents a potent initial transformative event towards chordoma formation.

## Methods

### Animal husbandry

Zebrafish (*Danio rerio*) were maintained, collected, and staged principally as described (*25*) and in agreement with procedures mandated by the veterinary office of UZH and the Canton of Zürich. If not indicated otherwise, embryos up to 5 dpf were raised in temperature-controlled incubators without light cycle at 28°C.

### Vectors and transgenic lines

*col2a1aR2:KalTA4 driver line:* The plasmid *pE5’-col2a1aR2_minprom* was cloned by amplifying the previously described *col2a1aR2 cis*-regulatory element (*26*) with the primers *col2a1a R2 fwd* and *col2a1a R2 BamHI reverse* from zebrafish genomic DNA using the Expand High Fidelity Polymerase System (Roche). The resulting PCR product was TA-cloned into *pENTR5’ (*Thermo Scientific) following the manufacturer’s instructions. The mouse *beta-Globin* minimal promoter (*minprom*) was amplified from *pME-minprom-EGFP (27*) with the primers *Mouse beta-Globin minimal promoter BamHI forward* and *Mouse beta-Globin minimal promoter XbaI reverse* and cloned into *pENTR5’-col2a1aR2*. The final plasmid *col2a1aR2:KalTA4,myl7:EGFP* was assembled by Multisite Gateway Cloning (see Supplementary Methods for details on cloning, transgenesis, and additionally used transgenic lines).

### Imaging and staining

Injected embryos were sorted by fluorescent dissecting scope-detectable notochord fluorescence (indicating the presence of the *UAS*-transgene) and imaged at 5 dpf. Animals with strong fluorescence were fixed in 4% PFA at 4°C overnight. Subsequently, embryos were embedded in paraffin, sectioned into 5 µm sections, deparaffinated, and stained with Hematoxylin and Eosin according to standard protocols. Immunohistochemical studies were performed according to the manufacturers’ protocol of the individual antibodies used (see Supplementary Methods for antibody details). Work on chordoma sections was performed under approval by BASEC-no. 2017-00017 by the Cantonal Ethics Commission Zürich. Live animals were imaged with a Leica M205FA stereo microscope. Histological sections were imaged with a Zeiss Axioscan Z1 Slidescanner using a Plan Apochromat 20x objective.

### Transmission Electron Microscopy

Wildtype and *HRAS*^*V12*^-expressing larvae where fixed 2.5% Glutaraldehyde at 8 dpf and processed for TEM at the Center for Microscopy and Image Analysis (ZMB), following standard protocols. Images were acquired with a Philips CM100 transmission electron microscope.

### Notochord isolation and mRNA-Seq

8 dpf *Tg(twhh:Gal4)* and *Tg(twhh:Gal4);Tg(UAS:EGFP-HRAS*^*V12*^) were euthanized with 3% Tricaine methanesulfonate (Sigma). Embryos were decapitated and incubated in Trypsin-EDTA (Sigma) for 30 minutes to facilitate tissue dissociation. Notochords were then dissected using tungsten needles and immediately transferred to Trizol-LS (Ambion). We isolated 30-50 notochords per replicate, with a total of 3 replicates per condition (3x wildtype, 3x *HRAS*^*V12*^). Notochord RNA was extracted following the manufacturer’s protocol using Trizol-LS. RNA-seq analysis was performed as outlined in Supplementary Methods.

## Results

### Sheath cell-directed oncogene expression drives chordoma formation in zebrafish

We first sought to functionally assess chordoma-associated genes that could initiate notochord hyperplasia in zebrafish embryos. While our previous chordoma modelling used as transgene driver *twhh:Gal4*, based on the *twhh* promoter region with promiscuous transcriptional activity in notochord cells (*24, 28*), we aimed for a stronger driver that would also enable injection-based oncogene testing. We generated *col2a1aR2:KalTA4* transgenics that base on the *R2* fragment of the *col2a1a* regulatory region (*26*) to express the optimized Gal4-based transactivator KalTA4 (*28*) (Fig. 1A,B). When incrossed with stable *UAS* reporters, *col2a1aR2:KalTA4* drove notochord-specific reporter expression detectable from 5-7 somite stage (approximately 12 hpf) in the entire notochord including sheath cells, expression which remained strong throughout development (Fig. 1A,B). *col2a1aR2* also drove *KalTA4* expression in other cartilage lineages, including the otic vesicle, jaw, and pectoral fin starting from 2.5 to 3 dpf throughout adulthood (Supplementary Fig. S1A-C), in agreement with previous analyses of *R2* activity (*26*).

**Figure 1:**
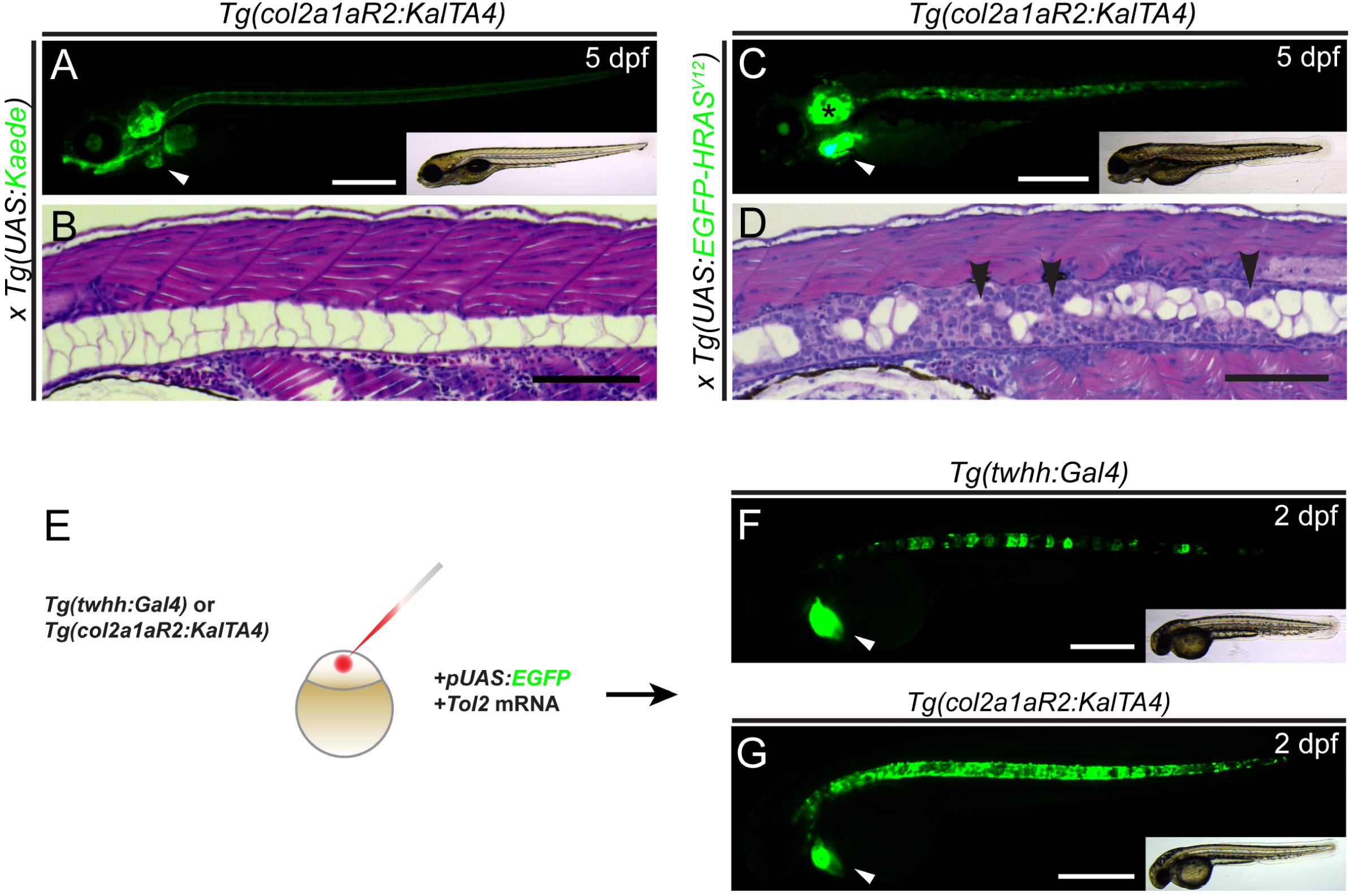
*Tg(col2a1aR2:KalTA4)* enables stable and transient oncogene expression in the developing notochord. (**A,B**) Lateral view (**A**) and transverse histology section (stained with Hematoxylin and Eosin (H&E), **B**) of 5 dpf zebrafish embryo transgenic for the transgene *col2a1aR2:KalTA4* and crossed to stable *UAS:Kaede (*visible as green fluorescence in **A**). Note expression in the notochord, craniofacial cartilage, otic vesicle, and pectoral fins; the prominent green heart (white arrowhead, **A**) indicates the *myl7:EGFP* transgenesis marker associated with *col2a1aR2:KalTA4*. The developing notochord shows the large vacuolated cells take up the vast majority of the notochord volume and are rimmed by a thin layer of sheath cells (**B**). (**C**,**D**) Expression of stable *UAS:EGFP-HRASV12 (*detectable by green fluorescence of the fusion protein, **C**) by *col2a1aR2:KalTA4* causes invasive and wide-spread notochord hyperplasia (black arrowheads, **D**) and overgrowth of other cartilage tissue (i.e. otic vesicle, asterisks in **C**); white arrowhead indicates *myl7:EGFP* transgenesis marker (**A**). (**E-G**) *col2a1aR2* provides a potent driver for transient notochord expression. (**E**) Injection of *UAS:EGFP* into the one-cell stage embryos with either *twhh:Gal4* or *col2a1aR2:KalTA4* to read out the resulting notochord mosaicism resulting from random integration of *UAS:EGFP* by Tol2 transposase. While injections into twhh:Gal4 results in highly patchy EGFP expression (**F**, green fluorescence, n=33/56), col2a1aR2:KalTA4 more consistently drives EGFP expression throughout the notochord (**G**, green fluorescence, 25/47); n indicates representative EGFP-expressing embryos in a injected representative clutch. Scale bars **A,C,F,G** =500 μm, **B,D**=200 μm. See also Supplementary Figure S1.

When we crossed *col2a1aR2:KalTA4* to stable *UAS:EGFP-HRAS*^*V12*^, double-transgenic embryos displayed over-proliferation of notochord sheath cells within 2 dpf: sheath cell proliferation advanced over time towards the center of the notochord, effectively compressing the inner vacuoles and excluding them altogether in parts of the notochord (Fig. 1C,D). This phenotype was similar, if not stronger, than the *twhh:Gal4;UAS:EGFP-HRAS*^*V12*^ combination used previously for chordoma modeling in zebrafish (*24*). In addition to the uncontrolled proliferation of notochord cells, *col2a1aR2:KalTA4;UAS:EGFP-HRAS*^*V12*^ embryos developed enlarged otic vesicles and deformed jaw cartilage starting from 4 dpf (Fig. 1C), consistent with the overall expression pattern of the *col2a1aR2:KalTA4* driver (Fig. 1A,C, Supplementary Fig. S1A-C). Taken together, these observations establish *col2a1aR2:KalTA4* as potent driver for notochord-specific transgene expression and notochord hyperplasia in zebrafish.

### *T/Brachyury* overexpression is insufficient to initiate notochord hyperplasia

Based on these results, we assessed if we can perform candidate gene tests by transforming notochord cells in transient injections into *col2a1aR2:KalTA4* embryos (Fig. 2A). Closely resembling the phenotype of stable *HRAS*^*V12*^ overexpression (Fig. 1C,D), transient *HRAS*^*V12*^ overexpression consistently caused localized tumorigenic lesions along the notochord (Fig. 2B-E). Corresponding to the mosaic pattern of the transient *HRAS*^*V12*^ expression resulting from UAS construct injection (Fig. 1E-G) and with stable transgenic expression (Fig. 1C,D), the transformed sheath cells locally invaded the notochord (Fig. 2E).

**Figure 2:**
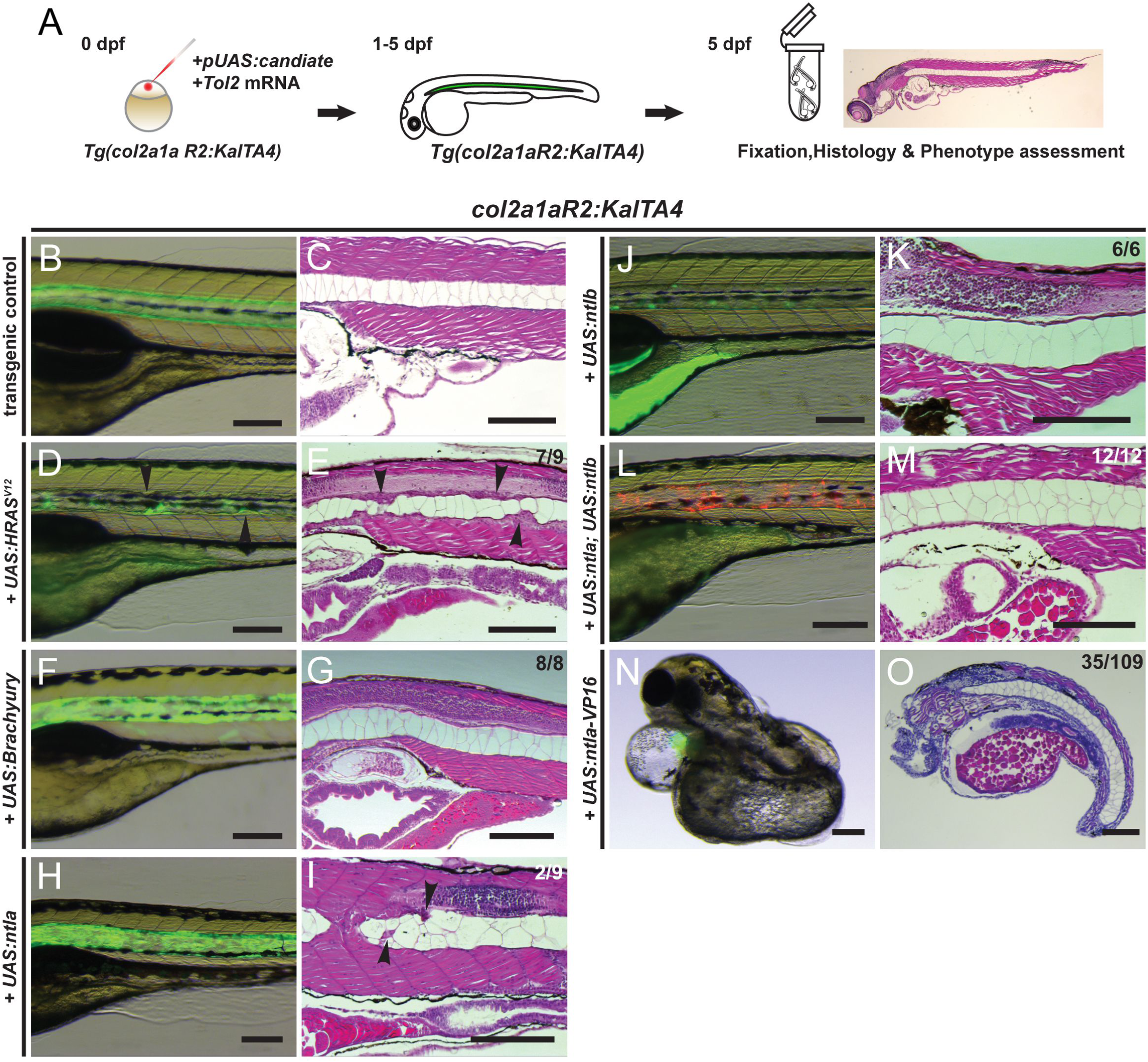
Overexpression of *Brachyury* genes in the zebrafish notochord is insufficient to initiate chordoma. (**A**) Workflow of injection-based notochord hyperplasia assessment: at the one-cell stage, *Tg(col2a1aR2:KalTA4)* embryos are injected with *Tol2* transposase mRNA and a plasmid containing a fluorescently-labeled candidate gene under *UAS* control; injected embryos are raised up to 5 dpf and candidate gene expression is monitored via notochord fluorescence. Embryos with consistent reporter expression are fixed, sectioned, and stained with H&E to assess the notochord phenotype with light microscopy. (**B-O**) Close-up lateral view of embryo notochords at 5 dpf, for brightfield and fluorescence (left column) and H&E histology (different embryos per condition); numbers indicate observed versus total from an individual representative experiment. (**B,C**) *col2a1aR2:KalTA4 c*ontrol reference at 5 dpf, expressing *UAS:Kaede* to fluorescently label the notochord. (**D,E**) Transient injection of *UAS:EGFP-HRASV12* causes localized notochord hyperplasia (arrowheads, **E**). (**F-I**) Forced expression of human *Brachyury (***F,G**) or of the zebrafish Brachyury gene *ntla (***H,I**) does not affect notochord development or cell proliferation. Minor lesions caused by collapsed vacuolated cells developed in few *UAS:ntla*-expressing notochords (arrowheads, **I**). (**J,K**) Overexpression of the second zebrafish Brachyury gene *ntlb* does not affect notochord development or proliferation. (**L,M**) Combined overexpression of both zebrafish *Brachyury* genes *ntla* and *ntlb* has no effect on proliferation and the notochord develops normally. (**N,O**) Notochord-driven expression of *ntla-VP16*, encoding a dominant-active transcriptional activator, leads to non-autonomous defects in the trunk with a shortened and/or severely curled trunk (32% of analyzed embryos), while the notochord forms normally. Scale bars 200 μm. See also Supplementary Figure S2.

We next applied *col2a1aR2:KalTA4* to test the potential of increased *Brachyury* activity to transform the developing notochord. We cloned *UAS* vectors harboring human *Brachyury* and its main zebrafish ortholog *ntla* coupled *in cis* with *UAS:EGFP* to avoid possible inactivating tagging of *Brachyury* proteins (*UAS:Brachyury,UAS:EGFP*, and *UAS:ntla,UAS:EGFP*, respectively). We chose this strategy as direct tagging of *Brachyury* ORFs with fluorescent proteins led to inconsistent transgene expression, possibly indicating aberrant protein folding or activity (Supplementary Fig. S2A-F). In contrast, upon injection of untagged *UAS-Brachyury* or *UAS-ntla* into *col2a1aR2:KalTA4* embryos, we observed reproducible *EGFP* fluorescence signal throughout the notochord, and all embryos expressing either human *Brachyury* or zebrafish *ntla* developed normal-appearing notochords within the observed 5 dpf (Fig. 2F-I). Consistent with the injection-based results, embryos carrying a stable *UAS:ntla* transgene and *col2a1aR2:KalTA4* also developed normal notochords (Supplementary Fig. S2G,H). Contrary to humans and mice that harbor a single copy of the *T/Brachyury* gene, zebrafish harbor two paralogs, *ntla* and *ntlb*. Nonetheless, the overexpression of *UAS:ntlb* alone did not affect the integrity of the notochord (Fig. 2J,K), nor did the combined overexpression of *UAS:ntla* and *UAS:ntlb* within the observed 5 days (Fig. 2L,M).

To augment the activity of zebrafish *ntla* as transcriptional activator, we expressed a Ntla-VP16 fusion protein wherein native Ntla is fused with the strong VP16 transactivation domain (*29*). *col2a1aR2:KalTA4*-driven notochord expression of transiently injected *UAS:ntla-VP16* reproducibly resulted in embryos with severe body curvature, a shortened body axis, and cardiac edema, yet analyzed notochords remained structurally intact and devoid of any hyperplasia (Fig. 2N,O).

Altogether, from these results within the timeframe of our observations, we conclude that increased expression of *ntl/Brachyury*, even of a transcriptionally hyperactive form, is *per se* insufficient to transform developing zebrafish notochord cells into a hyperplastic state.

### Overexpression of RTK genes found activated in human chordoma is sufficient to initiate notochord hyperplasia

Receptor tyrosine kinases (RTKs) and their branched downstream pathways are key players in development, and activation of RTKs and their downstream Ras-dependent cascades can transform a variety of cell types into tumors (*30*). While activated Ras mimics upstream RTK activation, as used in our original chordoma model (*24*), Ras mutations are seemingly rare in chordoma (*3, 31*). An increasing number of studies have rather found copy number alterations, increased phosphorylation, or misexpression of individual RTKs in chordoma, most prominently of *EGFR, KDR/VEGFR2, PDGFRA*, c-*KIT*, and different *FGFR* genes (*7*–*10*).

To test the potential of misexpressed individual RTKs for driving hyperplastic notochord transformation, we injected *col2a1aR2:KalTA4* embryos with *UAS* constructs harboring the ORFs of candidate RTKs and additional chordoma-implicated candidate genes as reference. Injection of *UAS* constructs for *c-KIT, PDGFRA, FGFR3*, and *FGFR4* all resulted in high mortality rates during somitogenesis (>80% in the case of *FGFR3 and FGFR4*): in injected embryos, we frequently observed severe body axis perturbations and dorso-ventral patterning defects (Supplementary Fig. S3A-D). These phenotypes suggest severe non-autonomous effects upon notochord-focused expression of *c-KIT, PDGFRA, FGFR3* and *FGFR4*, precluding further analysis using our approach. In contrast, and comparable to injection of *UAS-HRAS*^*V12*^ (Fig. 3A-D), notochord-driven expression of *UAS:EGFR* caused reproducible sheath cell hyperplasia between 2-5 dpf (Fig. 3E,F). Similarly, injection of *UAS*:*kdr/vegfr2* also potently triggered sheath cell hyperplasia (Fig. 3G,H). Both EGFR- and Kdr/Vegfr2-expressing notochords displayed localized hyperplasia along the entire length of the notochords (Fig. 3F,H): we repeatedly observed clusters of hyperproliferating sheath cells invading the center of the notochord, thereby compressing the vacuolated inner notochord cells, as also observed by *HRAS*^*V12*^ expression (Fig. 2E,3D).

**Figure 3:**
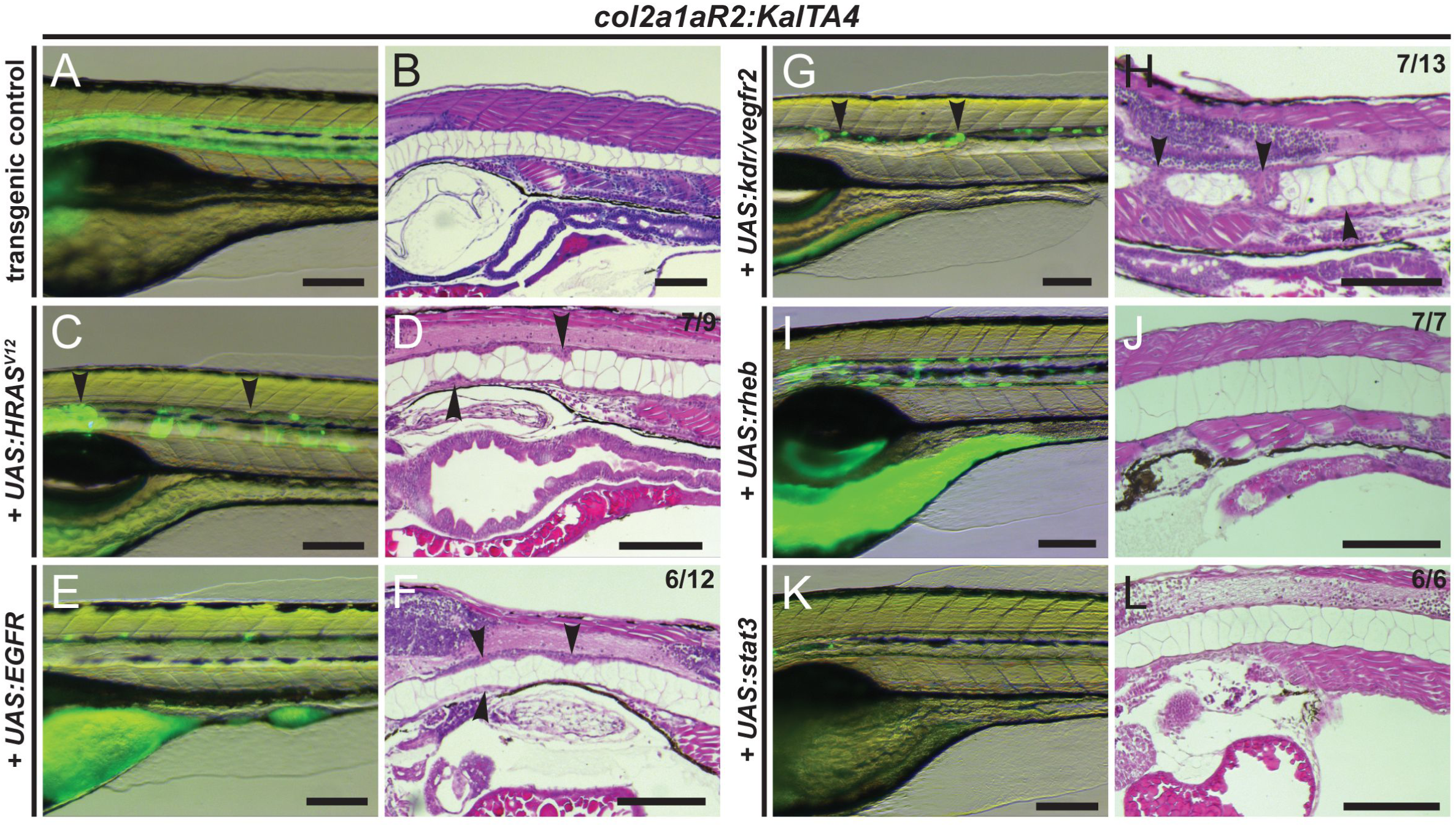
Transient overexpression of RTK genes drives notochord hyperplasia. (**A-L**) Close-up lateral view of embryo notochords at 5 dpf, with brightfield/fluorescence (left panels) and H&E histology (different embryos per condition; right panels); numbers indicate observed versus total from an individual representative experiment. (**A**,**B**) *col2a1aR2:KalTA4 c*ontrol reference at 5 dpf, expressing *UAS:Kaede* to fluorescently label the notochord. (**C,D**) Transient injection of *UAS:EGFP-HRAS*^*V12*^ causes localized hyperplasia (arrowheads, **C,D**) in the notochord. (**E,F**) Overexpression of human *EGFR* consistently causes local hyperplasia in the developing notochord (arrowheads, **F**). (**G,H**) Overexpression of zebrafish *kdr (*encoding the *VEGFR2* ortholog) causes strong hyperplasia (arrowheads in **H**, compare to **D**,**F**). (**I,J**) Zebrafish *rheb* overexpression leads to enlarged vacuoles and no detectable hyperplasia. (**K,L**) Zebrafish *stat3* overexpression leads to no detectable hyperplasia and allows for normal notochord development within 5 dpf. Scale bars 200 μm. See also Supplementary Figure S3.

Phosphorylation of mTOR, a key downstream mediator of RTK/Ras signaling in the mTORC1 and mTORC2 complexes in the control of ribosome biogenesis and protein synthesis, has been repeatedly found in chordoma (*32*). The small GTPase Rheb activates mTORC1 and can act upon overexpression as a proto-oncogene (*33, 34*). *col2a1a:KalTA4* embryos injected with *UAS:rheb* developed notably larger vacuolated notochord cells (Fig. 3I,J), in agreement with increased cell growth triggered by mTOR1 activity; nonetheless, the *UAS:rheb*-injected embryos did not develop any notochord hyperplasia within the observed 5 dpf (Fig. 3I,J). We further tested the tumorigenic potential of *STAT3* involved in JAK/STAT signaling which has been repeatedly found activated in chordoma (*31, 35*). Nonetheless, *col2a1aR2:KalTA4* embryos expressing *UAS:stat3* also developed normally without any signs of notochord hyperplasia (Fig. 3K,L). Taken together, our results suggest that of the genes we tested in our assay, misexpression of chordoma-implicated EGFR and KDR/VEGFR2 RTKs is sufficient to trigger the onset of specific notochord hyperplasia.

We confirmed that *EGFR-* and *kdr/vegfr2*-misexpressing cells have activated MAPK signaling, as revealed by probing for the downstream effector pERK: compared to 5 dpf wildtype notochords that are devoid of pERK (Fig. 4A), *HRAS*^*V12*^, *EGFR*, and *kdr/vegfr2* misexpression resulted in pERK-positive notochord cells, including prominent staining in cells infiltrating the notochord (Fig. 4B-D). Compared to controls, the *HRAS*^*V12*^-, *EGFR-*, and *kdr/vegfr2-*overexpressing notochords also stained notably stronger for the common chordoma marker pan-Cytokeratin (Fig. 4F-I). We also observed staining for zebrafish Brachyury protein Ntla (Fig. 4K-O; see Supplementary Fig. S4A-G for human *Brachyury* antibody). Consistent with the absence of hyperplastic cells, combined overexpression of the zebrafish *Brachyury* genes *ntla* and *ntlb* did not change the staining for any of these markers (except for the misexpressed Ntla itself) compared to wildtype (Fig. 4E,J,O). These observations suggest that commonly used diagnostic markers for human chordoma stain positive in hyperplastic zebrafish notochords with activated RTK signaling.

**Figure 4:**
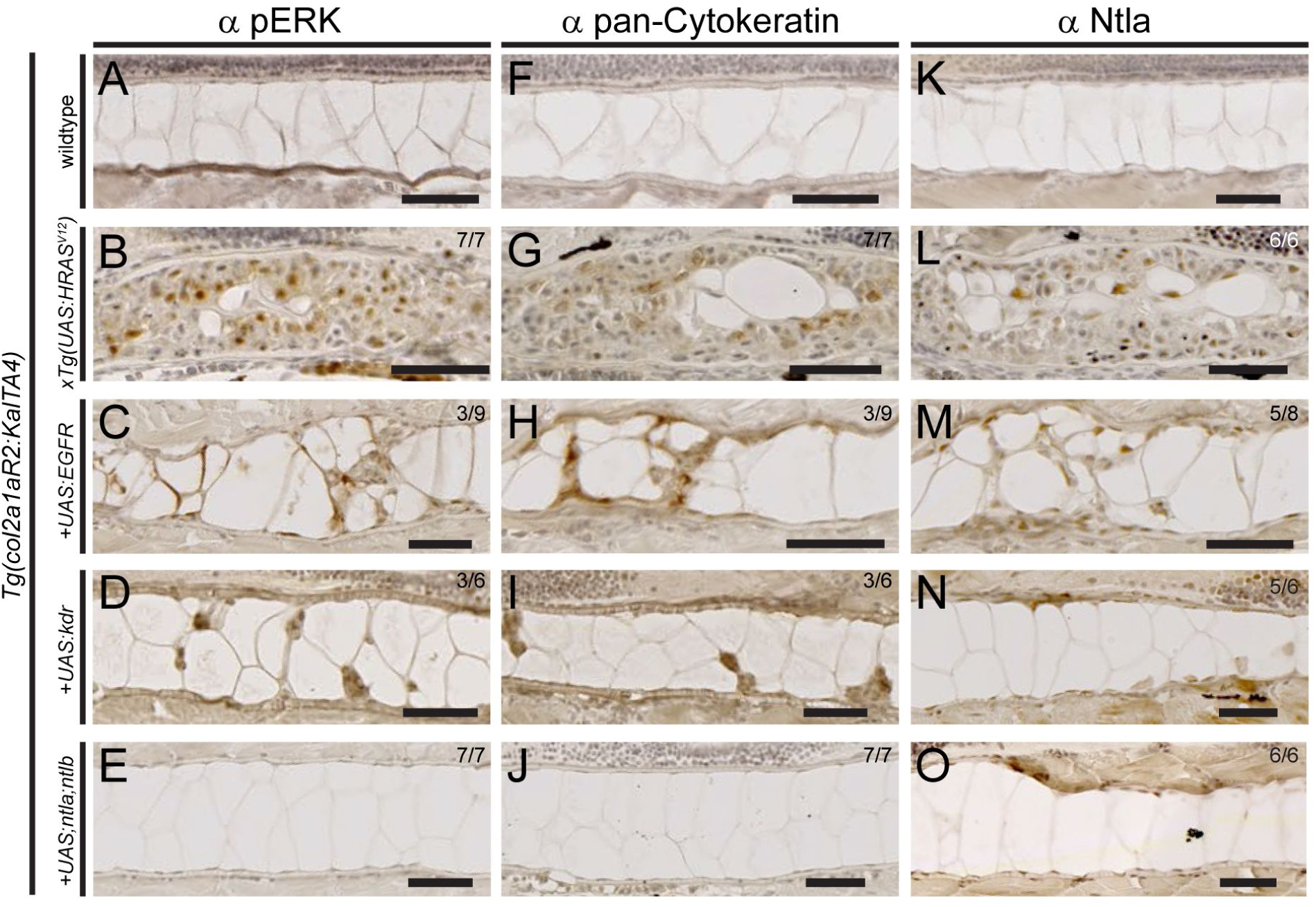
Expression of chordoma markers in RTK-transformed zebrafish notochords. (**A-O**) Immunohistochemistry on sagittal sections through the notochord of 5 dpf zebrafish embryos of the indicated genotypes, expressing either stable or mosaic transgenes. (**A-E**) MAPK pathway activation through *HRAS*^*V12*^, *EGFR*, and *kdr* overexpression results in nuclear pERK staining in the notochord, whereas controls and *ntla,ntlb-*injected embryos are negative for pERK staining. (**F-J**) *HRAS*^*V12*^, *EGFR*, and *kdr* oeverexpression results in staining for pan-Cytokeratin, whereas *ntla,ntlb-*injected embryos are negative for pERK staining akin to wildtype controls. (**K-O**) While control notochords show faint to no Ntla signal due to low cell density of the sheath layer (**K**), *HRAS*^*V12*^-overexpressing notochords (**L**) as well as in *EGFR-(***M**) and *kdr*-(**N**) overexpressing notochords show prominent nuclear Ntla staining. *ntla,ntlb*-overexpressing notochords also stain positive (**O**), confirming transgenic Ntla expression. Scale: 50 µm. See also Supplementary Figure S4.

Taken together, in zebrafish, increased levels of EGFR and KDR/VEGFR2 are sufficient to induce hyperplasia of developing notochord sheath cells by triggering RTK signaling, akin to constitutively active *HRAS*^*V12*^. In contrast, notochord-specific expression of effectors driving mTOR or JAK/STAT signaling, as also of different versions of *Brachyury (*Fig. 2F-O), did not trigger over-proliferation with early onset. These observations raise the possibility that aberrant activation of RTK signaling is a key process to trigger notochord cell hyperplasia, while other chordoma-implicated mechanisms could then potentially contribute after the initial tumor onset.

### Ras-mediated notochord transformation in zebrafish recapitulates abnormalities found in human chordoma

To gain more insight into the initial phase of notochord hyperplasia triggered by Ras-mediated notochord transformation in zebrafish, and to assess if genes associated with human chordoma become de-regulated, we performed RNA-seq analysis of dissected wildtype versus *HRAS*^*V12*^-transformed notochords (Fig. 5A).

**Figure 5:**
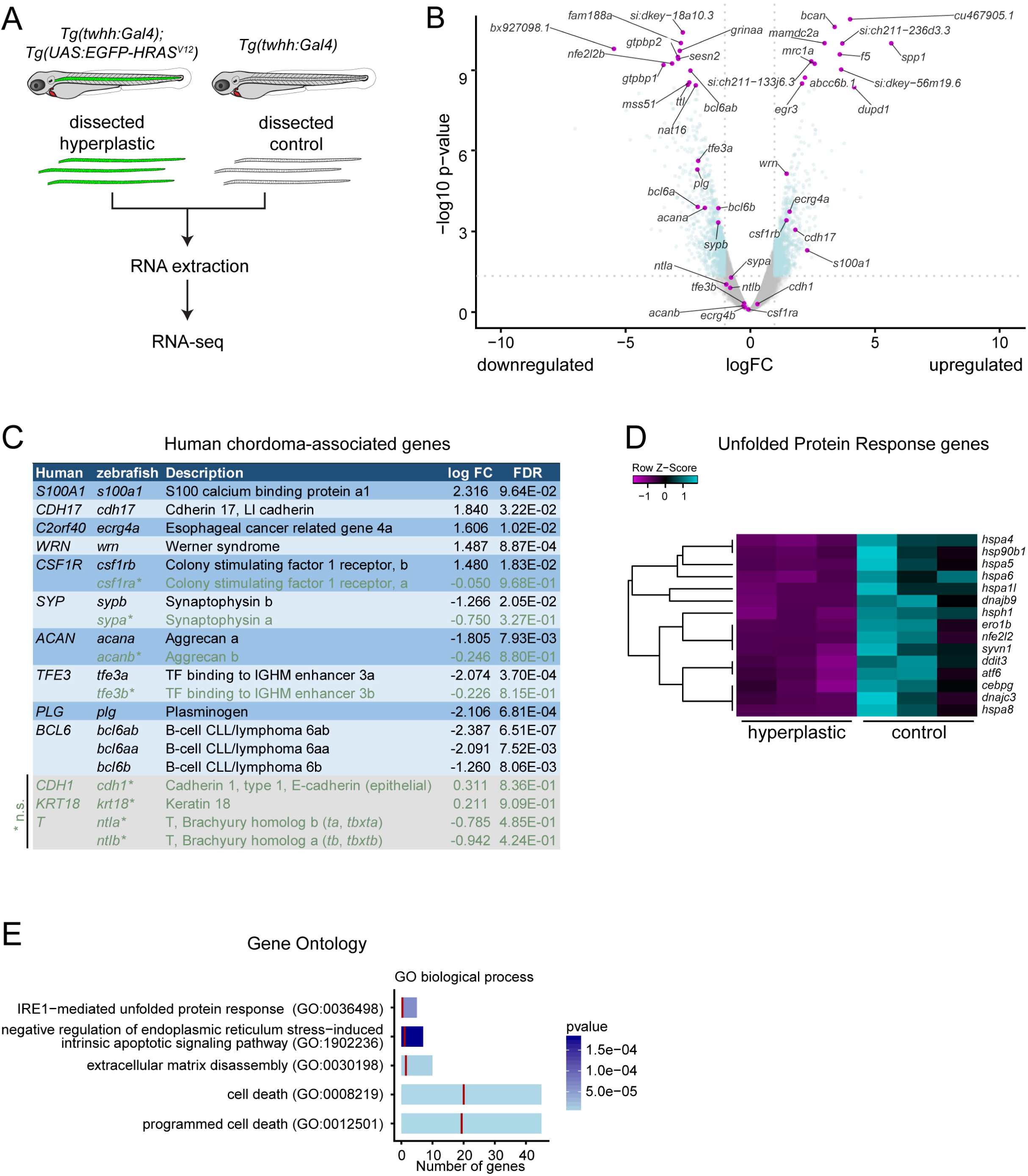
HRAS^V12^-induced zebrafish chordomas have a deregulated unfolded protein response and suppress apoptosis. (**A**) Workflow of control and transformed notochord isolation. Notochords were dissected from *twhh:Gal4 (*wildtype morphology and transgenic baseline) and *twhh:Gal4;UAS:EGFP-HRAS*^*V12*^-expressing larvae at 8 dpf; see Methods for details. (**B**) Volcano plot depicting overall distribution of de-regulated genes; grey indicating genes below p=0.05, turquoise indicating genes with significant de-regulation between control versus transformed notochords. Note that the zebrafish Brachyury orthologs *ntla* and *ntlb* are unchanged but expressed. (**C**) Expression of human chordoma-associated genes in HRASV12-transformed zebrafish notochords; see text for details. Zebrafish orthologs of human chordoma genes are shown with all their orthologs, with orthologs featuring not significant (n.s., green) changes marked with asterisks, genes with no significantly altered ortholog in grey boxes at bottom. (**D**) Significantly deregulated Unfolded Protein Response (URP) genes as per IPA pathway analysis of hyperplastic versus control zebrafish notochords. (**E**) Gene Ontology (GO) enrichment in hyperplastic zebrafish notochords also highlights deregulation of unfolded protein response, ER stress, ECM dynamics, and cell death. See also Supplementary Figure S5.

We found a total 591 genes as significantly deregulated in hyperplastic notochords compared to wildtype (228 upregulated, 363 downregulated; minimum log fold change ≥1, corrected FDR ≤0.05) (Fig. 5B, Supplementary Table S1). Notably, in the analyzed zebrafish control notochords, we detected modest but significant expression of both zebrafish *Brachyury*-coding genes *ntla* and *ntlb*, as confirmed by RT-PCR (Fig. 5B, S5A-C). Neither *ntla* nor *ntlb* significantly changed upon *HRAS*^*V12*^transformation (Supplementary Fig. S5A-C, Supplementary Table S1). These data indicate that, in the developing notochord, the expression of the zebrafish *Brachyury* genes *ntla* and *ntlb* persists beyond initial embryonic stages for considerably longer than commonly assumed based on mRNA *in situ* hybridization (*36*).

Among the significantly deregulated genes, we found genes that have previously been implicated in being aberrantly expressed in human chordoma. For instance, loss of the *BCL6* locus has been reported as possibly frequent event in chordoma (*6*). Consistent with this notion, all zebrafish *bcl6* genes, in particular *bcl6ab*, were significantly downregulated in our *HRASV12*-transformed zebrafish notochords (Fig. 5B,C, Supplementary Table S1). Further, *S100A1* is a potent diagnostic marker in clinical chordoma cases; zebrafish *s100a1* was also the most-significantly upregulated *s100* gene in our hyperplastic zebrafish notochords, albeit under slightly less stringent FDR threshold (logFC= 2.32, p=0.006, FDR=0.096) (Fig. 5B,C, Supplementary Table S1). We conclude that despite species differences, the induction of notochord hyperplasia in zebrafish using activation of the RTK/Ras cascade also deregulates genes observed in human chordoma.

We additional performed downstream pathway analysis of the deregulated genes using both Ingenuity Pathway Analysis and Gene Set Enrichment Analysis. Interestingly, both of these analyses revealed a significant downregulation of genes associated with ER stress and with the unfolded protein responses (Fig. 5D,E, Supplementary Fig. S5D-G). Given the established roles of UPR in notochord development and differentiation towards mineralized bone (*19*–*21*), this data suggests that one of the earliest responses to RTK/Ras transformation of the notochord is a suppression of this normal developmental program. Supporting this notion, other affected pathways included processes involved in bone and cartilage biology, including deregulation of markers of bone differentiation such as *spp1, anxa5*, and *ihh (*Supplementary Fig. S5E,G). In addition, we also noted a significant dysregulation of genes associated with extracellular matrix remodeling (Fig. 5E), a process associated with notochord sheath cells for building up a thick ECM around the forming notochord before the onset of segmented ossification (*26, 37*). Taken together, the transcriptome of notochord cells transformed by activated RTK/Ras signaling shows hallmarks of suppressed notochord differentiation.

To observe the cellular consequences of early notochord transformation as indicated by our transcriptome analysis, we performed electron microscopy (EM). Transverse sections analyzed by EM again documented the invasion of outer sheath cells towards the center of the notochord, resulting in compression of the inner vacuoles (Supplementary Fig. S6A,B). Strikingly, and in contrast to the small elliptical nuclei that occur in wildtype sheath cells, transformed notochord cells developed highly enlarged nuclei with irregular and lobulated shapes (Fig. 6A,B), as described previously (*38*). Notably, transformed notochord cells showed an increase in ER membranes throughout the analyzed transformed sheath cells (Fig. 6A,B), indicating highly active secretion that builds up unorganized ECM layers (Fig. 6C,D, Supplementary Fig. S6). Zebrafish chordomas generated with our approach do not metastasize (Burger et al., 2014; this study) but remain confined to the notochord. The ECM could, ultimately, be the cause for the absence of detectable outgrowth from the notochord in our zebrafish model: we observed several instances of individual sheath cells that had seemingly detached from the notochord and had become completely entombed within the secreted ECM (Fig. 6D). We detected a high abundance of secretory vesicles budding with the cell membrane in wildtype notochords, while transformed notochord cells featured amorphic cell boundaries to the ECM with trapped membrane fragments embedded in the collagen matrix (Fig. 6E,F).

**Figure 6:**
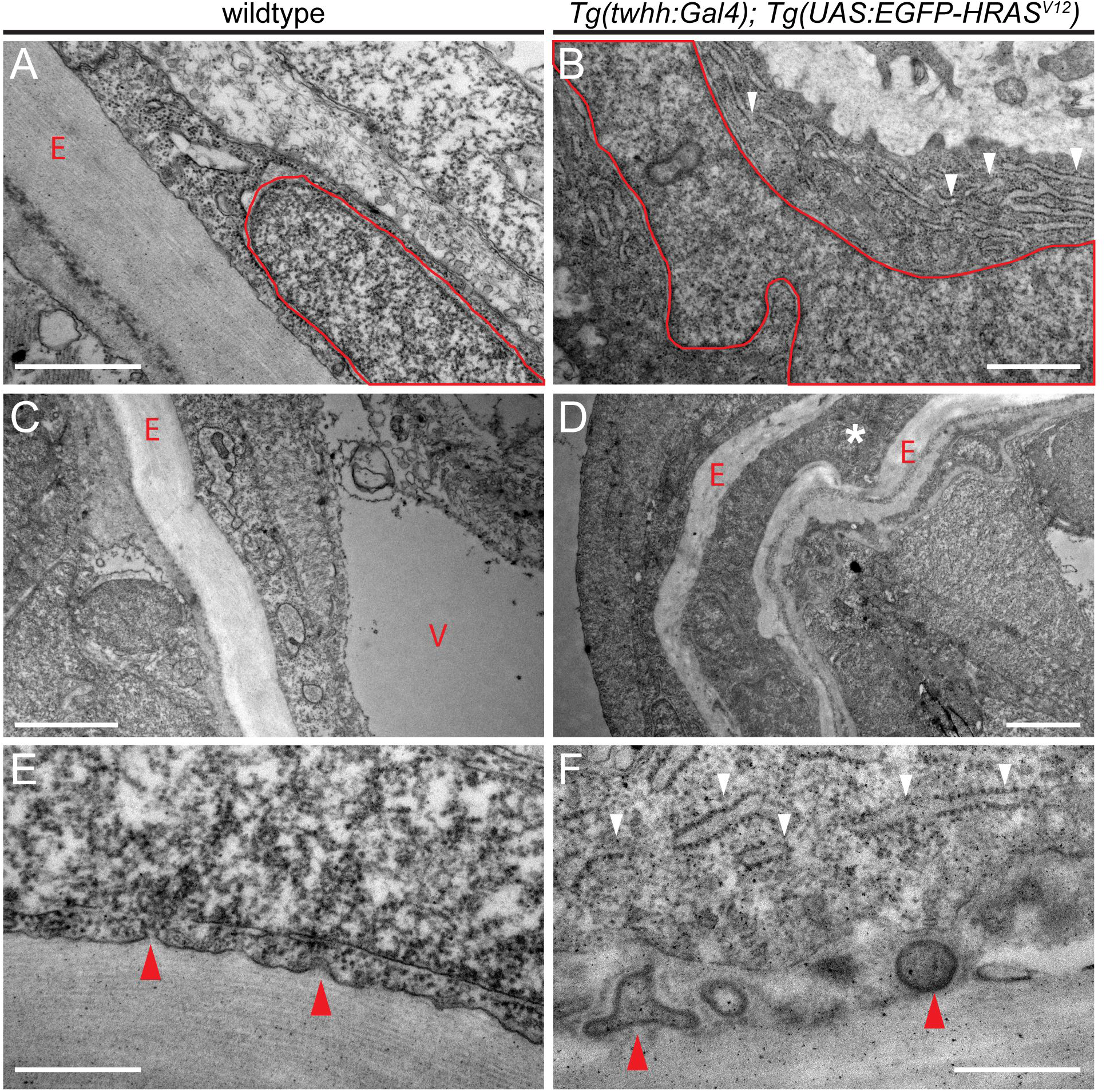
Aberrant ECM and ER accumulation in *HRAS*^*V12*^-induced zebrafish chordomas. (**A-F**) Transverse section through wildtype (**A**,**C**,**E**) and HRASV12-transformed zebrafish notochord (**B**,**D**,**F**) at 8 dpf imaged with transmission electron microscopy (EM). (**A**,**B**) Wildtype cells have a pill-shaped, regular nucleus (red outline, **A**) and form regular ECM layers (red E, **A**) secreted by the sheath cells at the outside of the notochord. Nuclei in transformed cells expand and develop lobed and distorted nuclear shapes (red outline, **B**); transformed cells further accumulate extensive ER lumen (white arrowheads, **B**). (**C**,**D**) In wildtype notochords, vacuolated cells (red v, **B**) take up the majority of the space inside the ECM-lined (red E, **C**) notochord to provide mechanical stability; transformed notochords become filled with non-vacuolated cells and secrete aberrant amounts of ECM that lead in extreme cases to entombed cells trapped in ECM layers (red asterisk, **D**). (**E,F**) Membrane details of wildtype versus transformed notochord sheath cells. Wildtype notochords show budding of vesicles that transport collagen for stereotypically layered ECM buildup (red arrowheads, **E**); in transformed notochords, the secretion process appears overactive (ER accumulation shown by white arrowheads, **F**) and results in membrane inclusions within the ECM (red arrowheads, **F**). See also Supplementary Figure S6.

Taken together, these observations suggest that RTK/Ras cascade-triggered sheath cell hyperplasia features hallmarks of deregulated ER stress and UPR. Oncogenic transformation by aberrantly activated RTK signaling could therefore drive the maintenance of an incompletely differentiated, developmental state in transformed notochord cells. In contrast, Brachyury expression is ongoing in the zebrafish notochord both in wildtype and in transformed conditions, indicating that maintained Brachyury expression reflects notochord identity of the transformed cells.

## Discussion

While assumed to represent a notochord-derived tumor, the exact ontogeny and hyperplasia-inducing events leading to chordoma remain unclear. To our knowledge, activating Ras mutations have so far not been reported from chordoma samples, and a comprehensive comparison of the zebrafish-based chordoma transcriptome with human chordoma is hindered by the lack of readily accessible transcriptome data of this rare tumor. Nevertheless, several RTK genes have been repeatedly found aberrantly activated or amplified in human chordoma samples. Harnessing our zebrafish-based notochord readouts as proxy for chordoma onset, we evaluated the potential of several chordoma-implicated candidate genes for their capacity to transform native notochord cells *in vivo*, both with injection and verified with stable transgenic insertions. In our assays, the RTKs EGFR and KDR/VEGFR2, both abundantly found as oncogenes in various other cancer types, robustly triggered notochord hyperplasia. In contrast, overexpression of various forms encoding *Brachyury*, including as hyperactive VP16 fusion protein, caused no apparent notochord transformation in our observed timeframe. These results provide the first direct, functional testing of the chordoma-inducing potential of implicated oncogenes in native notochord cells.

We performed overexpression of *Brachyury* in the developing notochord by different means to avoid use of functionally impaired fusion proteins (Supplementary Fig. S2A-E). Misexpression of human *Brachyury* as well as individual or combinatorial expression of the zebrafish *Brachyury* genes *ntla* and *ntlb (*most-recently renamed *tbxta* and *tbxtb*), consistently failed to induce notable notochord hyperplasia (Fig. 2F-M). Of note, their misexpression did occasionally cause aberrant notochord architecture, with collapsed sheath cells reminiscent of recent reports of structural lesions in notochord morphology (Fig. 2H,I) (*39*). Most compelling is the inability of *ntla-VP16* to transform the notochord: *ntla-VP16* encodes a previously validated (*29*), highly transcriptionally active fusion protein through the viral VP16 transactivation domain (Fig. 2N,O). These results provide first *in vivo* evidence that, in the zebrafish within the first 5 days, misexpression of Brachyury in developing notochord cells is insufficient to elicit a hyperplastic response.

Besides Brachyury, pathological detection of several other factors has been recurrently reported in chordoma cases. Ras/PI3K/AKT pathway activators, mainly RTKs including *EGFR* and *KDR/VEGFR2*, are frequently found activated or even genetically amplified in chordoma patients (*7*–*10, 32*). Further, promising advances in experimental chordoma treatment have been using RTK inhibitors, specifically compounds targeting *EGFR (40, 41*). Our results from assessing the hyperplasia-inducing potential of individual factors suggest that several RTKs are potent, and possibly redundant, oncogenes when aberrantly activated in notochord cells (Fig. 3E-H). While RTKs relay critical signals during embryo development, as underlined by the severe developmental defects caused by notochord-focused misexpression of *FGFR, Kit*, and *PDGFR* genes (Supplementary Fig. S3A-D), the repeatedly chordoma-associated *EGFR* and *KDR/VEGFR2* are, in our assay, individually sufficient to transform developmental notochord cells (Fig. 3E-H). Phosphorylated mTOR, the core component of mTORC1 and mTORC2 downstream of activated RTKs, was found in a number of chordoma cases (*4, 32, 42*). Nonetheless, activating mTORC1 via misexpression of its direct upstream regulator Rheb only results in morphological changes without hyperplasia (Fig. 3I,J), suggesting that additional events downstream of RTK/Ras activation are required for triggering notochord hyperplasia. *STAT3*-based signaling has been implicated in several chordoma studies (*43*), yet does not seem to be universally activated in chordoma. In our assay, misexpression of *STAT3* had no discernible impact on the notochord (Fig. 3K,L), suggesting the potential of *STAT3* to promote chordoma could contribute after the tumor-initiating hits.

Of note, our results with negative hyperplasia outcome do not rule out a chordoma-promoting potential for *Brachyury, STAT3*, or any other tested factor. Our assay is confined to an early developmental time window, which has the potential to reveal the sufficiency of aggressive notochord-transforming factors. Our results rather suggest that the chordoma-initiating events are most-potently mediated by triggering upstream events of RTK signaling; further work is warranted to analyze the synergy and combinatorial action of the individual chordoma lesions found in particular patients or across the analyzed tumors so far. In particular, the striking consistency of *Brachyury* expression in chordoma has gathered increasing attention as tumor-defining and possibly causative characteristic, pointing at the re-activation or maintenance of the embryonic notochord program as culprit (*12, 14, 44, 45*). Nonetheless, developmental perturbation of *Brachyury* during mouse notochord formation suggests a role in maintaining cellular notochord identity, without impact on proliferation or other cellular phenotypes upon perturbation (*17*). Further, the relevance of *Brachyury* expression for patient outcome remains unclear.

Two possibilities could account for the prominent expression of *Brachyury* in chordoma. First, in line with previous reports (*17, 27*), *Brachyury* expression reflects the notochord identity of transformed notochord remnants. Several upstream signaling pathways influence *Brachyury* expression during development by acting on *cis*-regulatory elements that remain incompletely charted. Consequently, *Brachyury* expression in transformed notochord cells might simply reflect the sustained activity of the developmental notochord program. This conclusion is further supported by our observation of persistent *ntla/ntlb* expression in zebrafish that did not significantly react to RTK/Ras activation (Fig. 5B, Supplementary Fig. S5B,C, Supplementary Table S1), and similar reports of sustained *Brachyury* expression in mouse (*17*) and human (*46*).

Alternatively, not acting as transforming agent by itself, *Brachyury* expression could feed into the aberrant transcriptional program ongoing in chordoma cells after initial transformation. In chordoma, concomitant elevated *Brachyury* expression could result in additional or combinatorial events that direct RTK-transformed cells into a hyperproliferative state conductive to tumor progression and metastasis formation. Congenital amplification of the *Brachyury* locus could confer sensitivity to notochord cells for aberrant RTK activation and for subsequently faster tumor formation (*14*). *Brachyury* knockdown in the developing notochord in mouse has revealed that its function was dispensable for progenitor cell survival, proliferation, and EMT (*17*). Expressed beyond physiological levels for a prolonged time, *Brachyury* could nonetheless contribute aberrantly to tumorigenic events. *Brachyury* can influence the RTK-based FGF pathway by controlling production of the FGF2 ligand to possibly maintain a positive feedback loop, resulting in a mesenchymal phenotype (*47, 48*).

Our RNA-seq analysis of Ras-transformed zebrafish notochords mimicking activated RTK signaling revealed deregulation of clinically relevant chordoma genes, including *s100a1* and *bcl6* family members (Fig. 5). These changes were concomitant with alterations in UPR, ER stress response, and ECM pathways (Fig. 5, Supplementary Fig. S5). Ultimately, the excessive accumulation of ECM collagen sheets around the transformed notochord likely hinders the hyperplastic cells in their expansion (Fig. 6). Hence, RTK-based transformation alone might skew notochord cells into a cellular state that caps their proliferative and invasive potential. Consistent with this notion, we detected deregulated UPR and ER stress pathways, which were accompanied by excessive ECM buildup that ultimately entombed individual cells (Fig. 6E,F). Coordinated activation of secretory pathway features with concomitant activation of the UPR is a hallmark of progressive notochord differentiation in medaka and Xenopus (*19, 21, 49*). The prominent secretory activity of differentiating notochord cells, in particular collagen secretion and ECM buildup, requires careful fine-tuning by the UPR to prevent apoptosis and aberrant protein accumulation in the ER (*50*). We therefore hypothesize that the suppression of an UPR and ER stress signature upon RTK/Ras activation in notochord cells prevents their terminal differentiation, keeping them in a proliferative, progenitor-like state that is susceptible to additional oncogenic insults. Therapeutic targeting of RTK signaling could consequently attack a main pathway required for the initial transformation of native notochord cells towards chordoma.

## Supporting information

## Acknowledgements

We thank Sibylle Burger and Seraina Bötschi for technical and husbandry support; Dr. Stephan Neuhauss for zebrafish support; the ZBM and Pathology Core of the Institute of Anatomy at UZH for imaging and sample prep support; Dr. Rodney Dale for critical input on generating the *col2a1a* transgenics; Dr. Ashley Bruce for the *ntla-VP16* construct; Dr. Claudia Palena for sharing the human *pBrachyury* vector; Dr. Andy Oates for the anti-Ntla antibody; and Craig Nicol, Alessandro Brombin, and Dr. Elizabeth Patton for sharing schematics. This work has been supported by an UZH URPP “Translational Cancer Research” seed grant to A.B.; a project grant from the Swiss Cancer League and the SwissBridge Award 2016 from the SwissBridge Foundation to C.M. and A.B.; a Swiss National Science Foundation (SNSF) professorship [PP00P3_139093], a Marie Curie Career Integration Grant from the European Commission [CIG PCIG14-GA-2013-631984], the Canton of Zürich, and the UZH Foundation for Research in Science and the Humanities to C.M; a UZH CanDoc fellowship to G.D.

## Author Contributions

G.D., C.M., and A.B. designed, performed, and analyzed the experiments; E.M.C., M.R., and R.M.W. performed transcriptome analysis and data interpretation; R.K. supported EM experiments; K.F. and E.J.R. provided the paraffin-embedded chordoma biopsies; C.M. and A.B. supervised the project; G.D., E.M.C., C.M., and A.B. compiled data and wrote the manuscript.

## Supplementary Figures

**Supplementary Figure S1:**
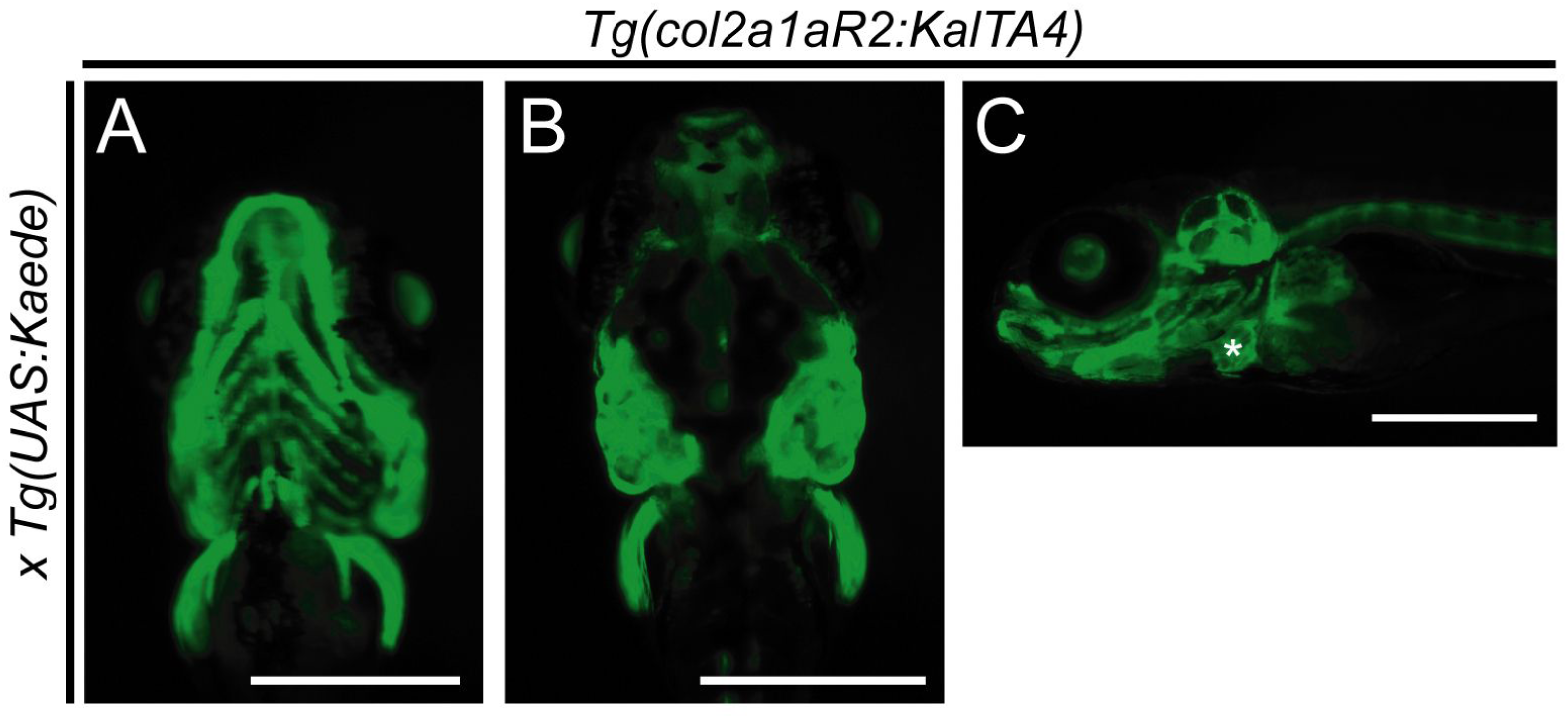
Activity of *Tg(col2a1a:KalTA4)* at later developmental stages and adults. (**A-C**) *col2a1aR2:Kal4TA4* activity shown by crossed-in *UAS:Kaede* expression (green fluorescence). (**A**) Ventral view showing the developing jaw cartilage and pectoral fins at 5 dpf. (**B**) Dorsal view showing activity in otic vesicle cartilage and pectoral fins at 5 dpf. (**C**) Lateral view of the craniofacial cartilage expressing *col2a1aR2:KalTA4* at 5 dpf. See also Figure 1.

**Supplementary Figure S2:**
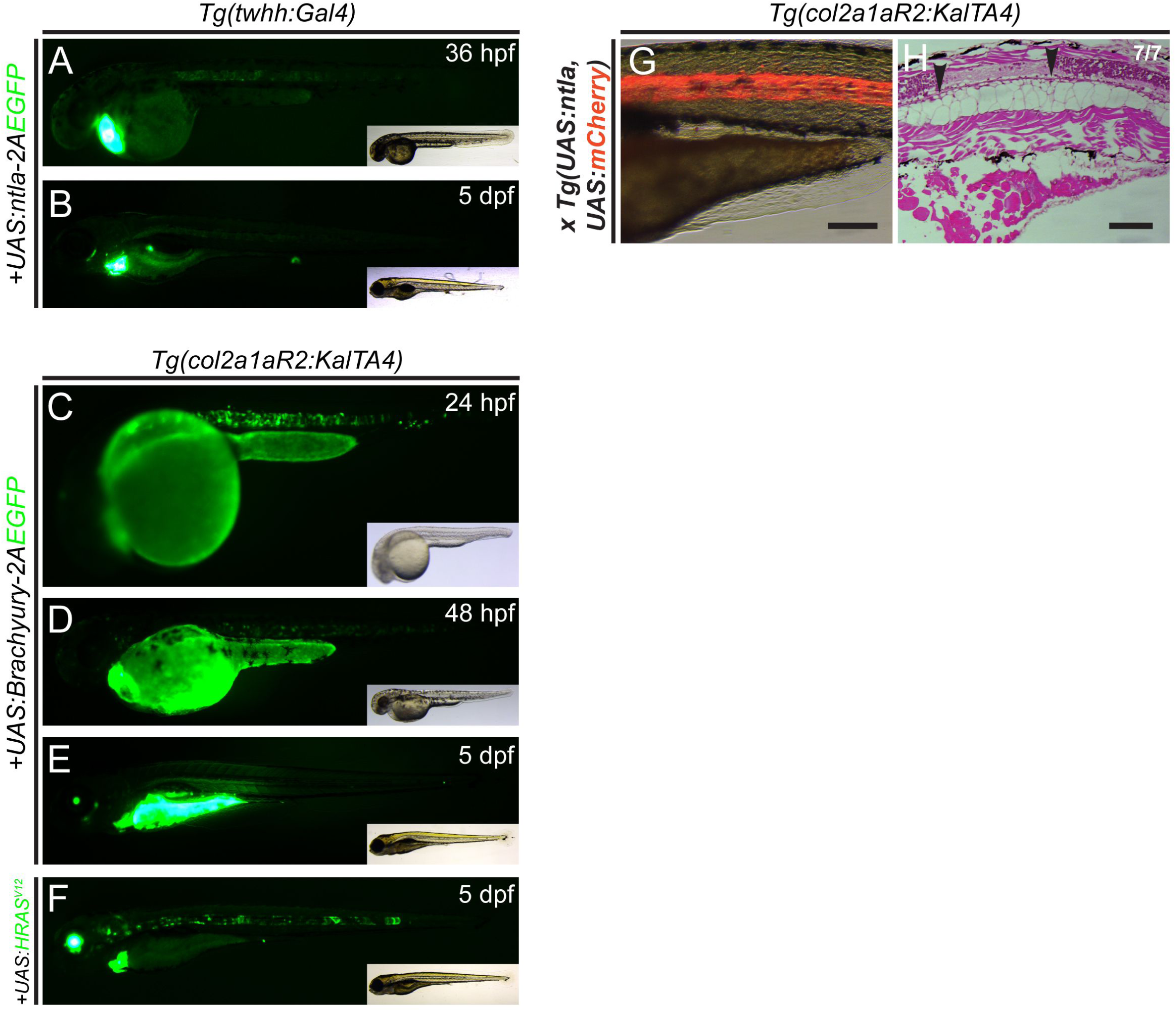
Tagged fusion proteins of Brachyury fail to express efficiently *in vivo*. (**A-F**) Lateral views of zebrafish embryos at indicated developmental stages, with EGFP fluorescence (green) in main panels, insets show brightfield for overall embryo morphology. (**A**,**B**) Transgene silencing in *Tg(twhh:Gal4)* with injected *UAS:ntla-2A-EGFP*; while expressed specifically in the notochord at 36 hpf (n= 43/56), expression greatly reduced at 5 dpf. (**C-F**) Representative embryos transiently expressing *UAS: Brachyury-2A-EGFP* at 24 hpf at detectable levels (**C**), while expression becomes significantly reduced at 48 hpf (**D**) and is completely silenced at 5 dpf (**E**). (**F**) Transiently expressed *UAS:EGFPHRAS*^*V12*^ at 5 dpf mainly localizes in the notochord and to a minor extent in the otic vesicle. Tumorigenic lesions could only be detected in the notochord. Transgenic markers used are *myl7:EGFP* for *twhh:Gal4* and *col2a1aR2:KalTA4, cryaa:Venus* for injected *UAS-*plasmids. (**G**,**H**) Akin to injected *UAS:ntla*, stable transgenic *Tg(UAS:ntla) o*verexpression fails to cause any significant notochord defect, with only few areas showing smaller vacuolated cells (arrowheads, **H**). See also Figure 2.

**Supplementary Figure S3:**
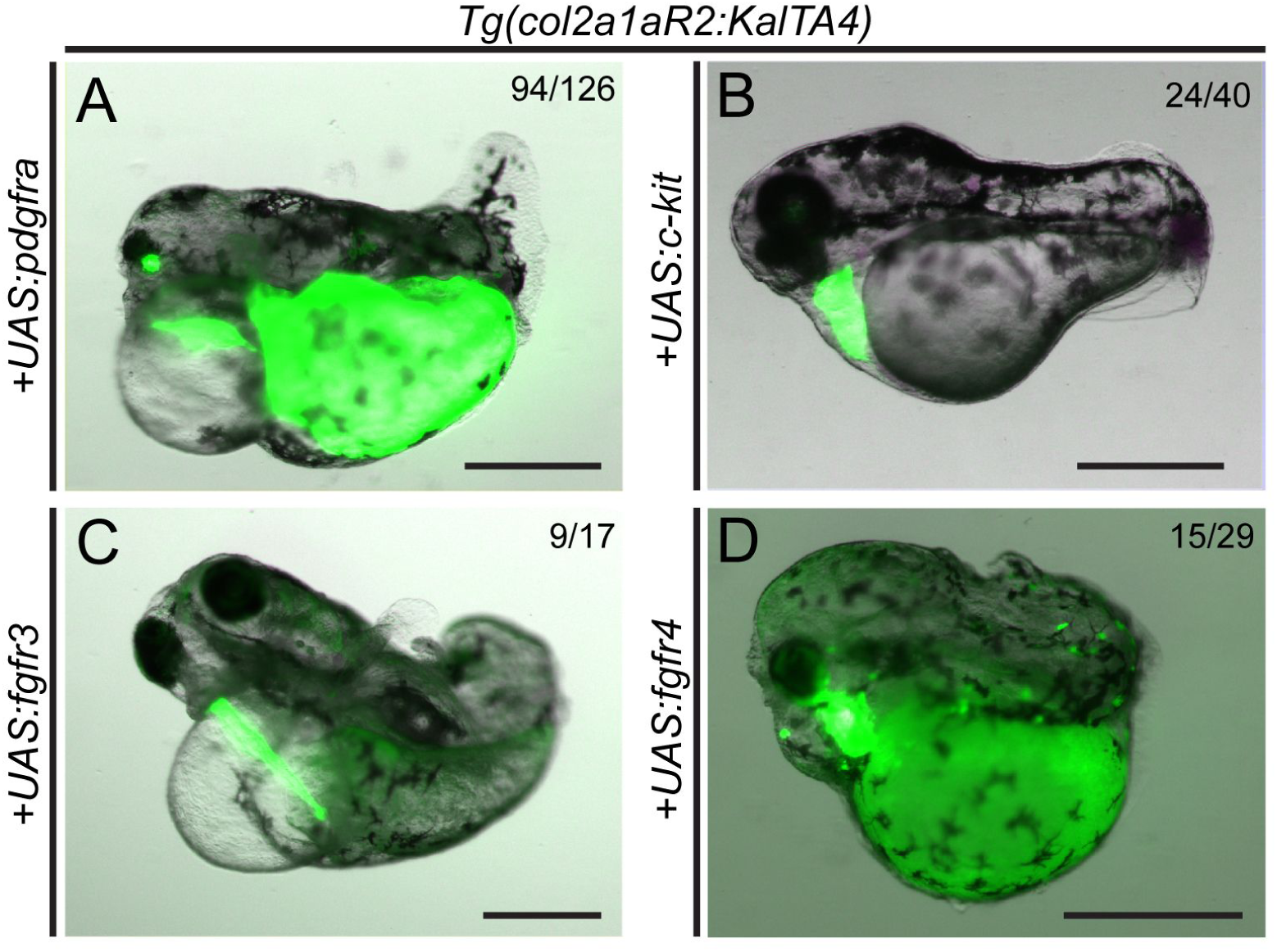
Embryo-wide, non-autonomous perturbation of embryo development upon notochord-specific overexpression of individual developmental RTK genes. (**A-D**) Representative *Tg(col2a1aR2:KalTA4)* embryos injected with individual *UAS* constructs for different chordoma-implicated RTK genes; numbers depict observed phenotype per total embryos in a representative experiment. Notochord specific overexpression of zebrafish *pdgfra (***A**), *c-kit (***B**), *fgfr3 (***C**), and *fgfr4 (***D**) leads to severe body axis truncation and aberrant development of cardiovascular and craniofacial structures (**A, C, D**: 5 dpf, **B**: 3 dpf). Transgenic markers: *myl7:EGFP* for *col2a1aR2:KalTA4, cryaa:Venus* for injected UAS plasmids. Scale: 500 µm. See also Figure 3.

**Supplementary Figure S4:**
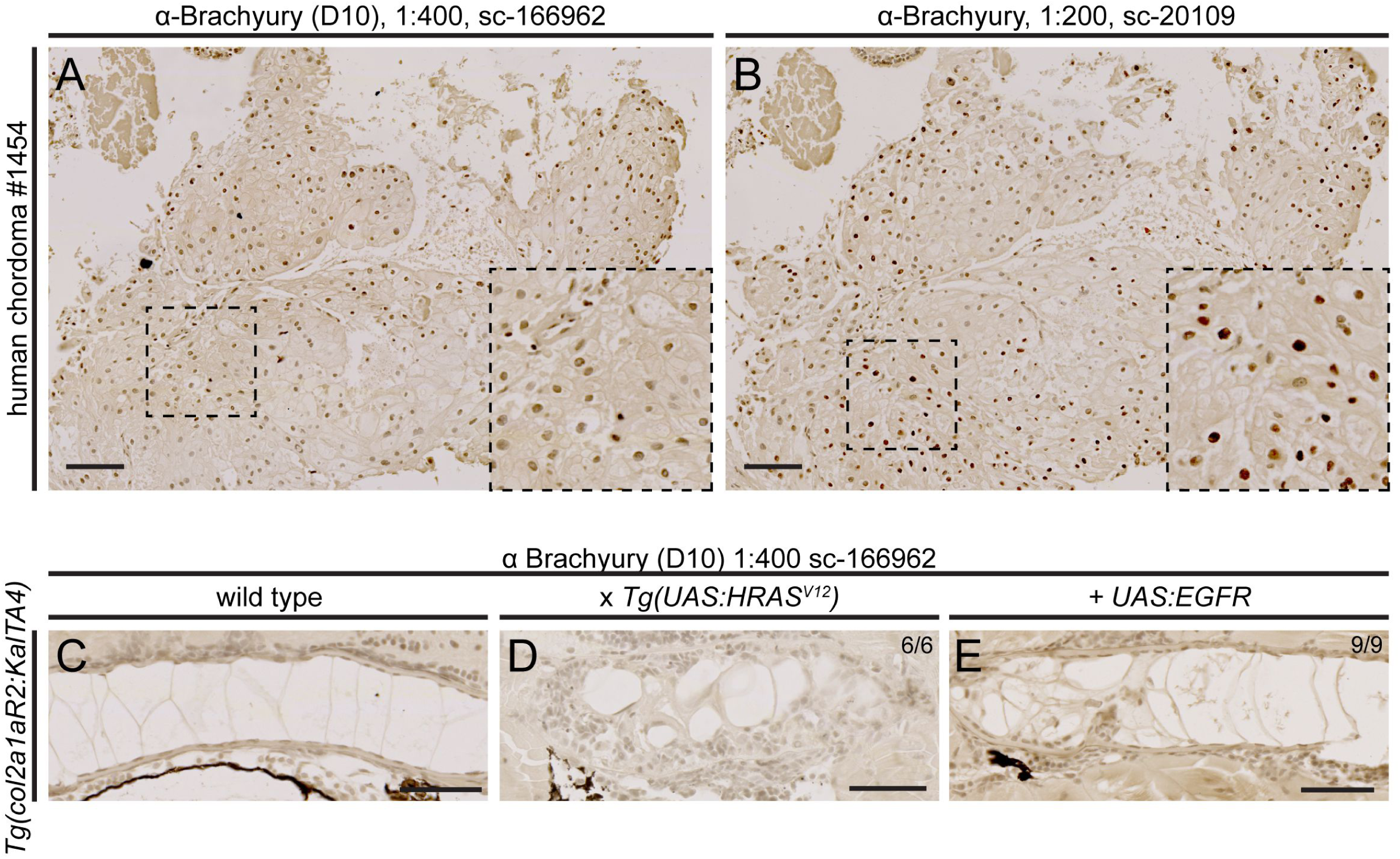
Testing of anti-Brachyury antibodies on human chordoma samples and zebrafish notochord sections. (**A,B**) Human chordoma sections from same tumor (chordoma #1454), stained for Brachyury using α-Brachyury D10 (sc-166962) and a discontinued α-Brachyury antibody (sc-20109) as positive staining control. Both reagents produce a strong nuclear staining. (**C-G**) α-Brachyury D10 fails to stain zebrafish tissue in several overexpression scenarios. See also Figure 4.

**Supplementary Figure S5:**
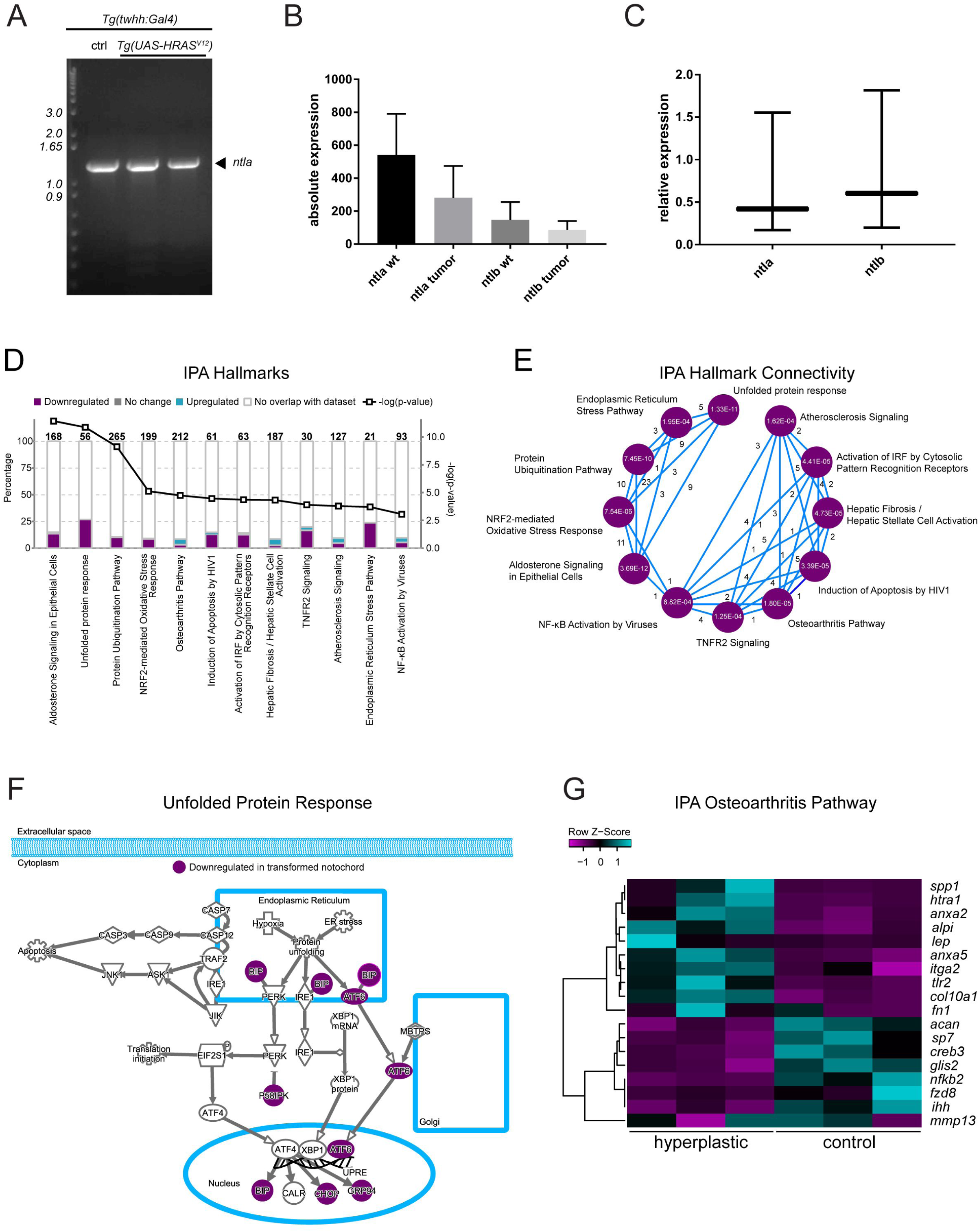
Transcriptome analysis of wildtype versus transformed zebrafish notochords reveals maintained expression of *Brachyury* genes and deregulated pathways. (**A-C**) Zebrafish *Brachyury* genes *ntla* and *ntlb* genes remain expressed beyond early development. (**A**) RT-PCR of isolated 8 dpf wildtype and *HRAS*^*V12*^-expressing notochords (two independent samples), respectively, confirms ongoing expression of *ntla*. (**B**,**C**) Absolute (number of reads) and relative levels of *ntla* and *ntlb* expression in wildtype and *HRAS*^*V12*^-expressing notochords. Compared to *ntla, ntlb* is also expressed but at lower levels. Differences between wildtype versus *HRAS*^*V12*^-expressing samples are not significant. (**D**) Ingenuity Pathway Analyis (IPA) output for the twelve most-significantly enriched pathways deregulated in transformed notochords; note repeated association with protein stress, unfolded protein response (UPR), and secretion of the enriched pathways. (**D**) Circle plot of main uncovered processes via IPA, with blue lines connecting Hallmark sets that share genes (numbers associated with blue lines); numbers inside Hallmark circles represent p-values. (**E**) Schematic of the unfolded protein response (URP) pathway, with components found downregulated in transformed zebrafish notochords shown in color. (**G**) Expression heat map of the IPA hallmark Osteoarthritis Pathway, encompassing key regulators of bone formation that are de-regulated between control versus hyperplastic zebrafish notochords. See also Figure 5.

**Supplementary Figure S6:**
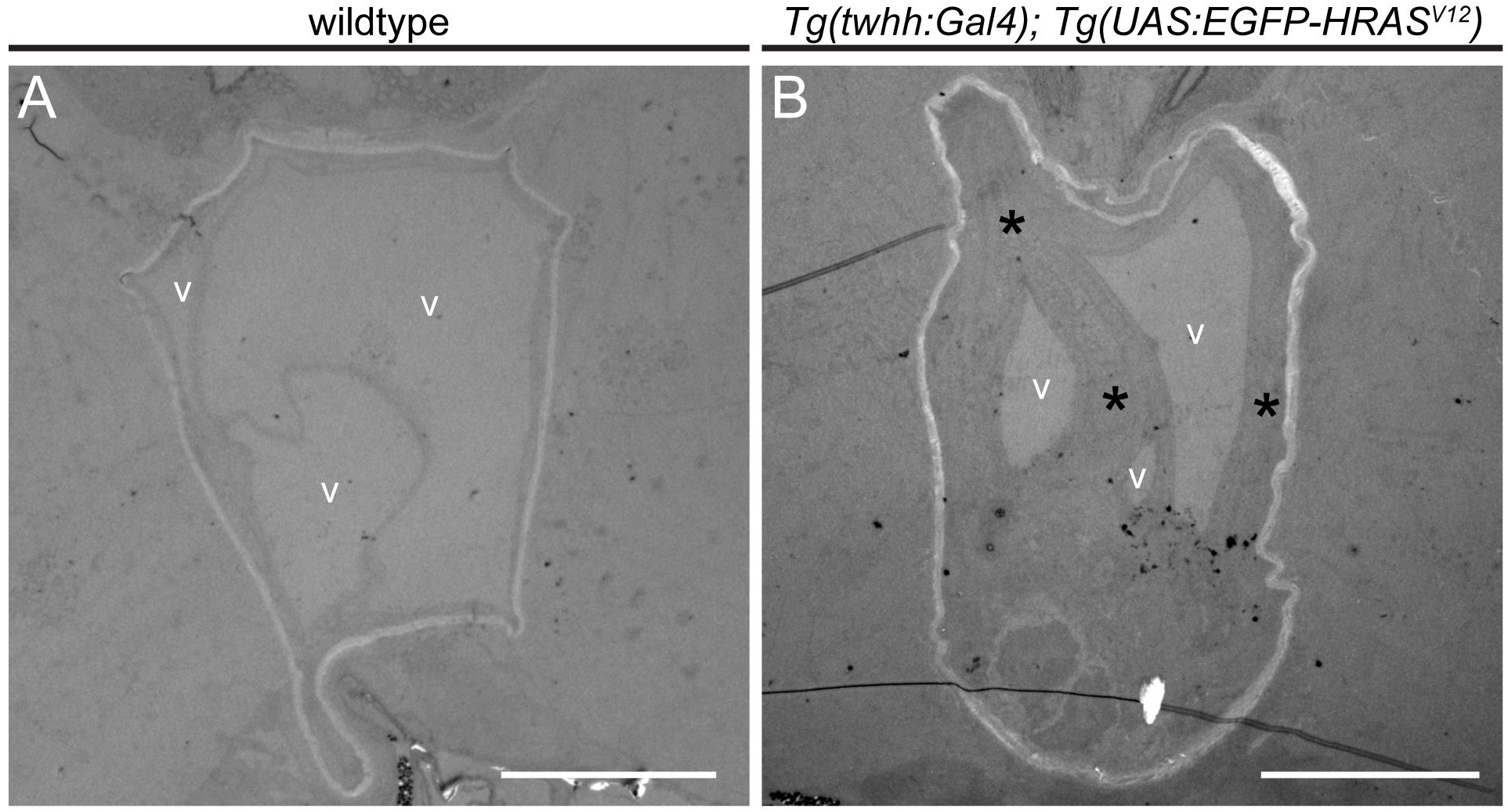
Overview of notochord sections used for EM. (**A,B**) Transverse section through a wildtype (**A**) and a representative *HRAS*^*V12*^-expressing notochord (**B**). Vacuolated cells (V) make up the majority of the notochord volume in wildtype, but are are markedly smaller upon transformation due to overproliferating sheath cells converging towards the center of the notochord (asterisks in **B**). Note the light band of ECM surrounding the notochords. Deformation of the natively circular notochord shape results from fixation procedures for EM. See also Figure 6.

**Supplementary Table S1: RNA-seq results** Compiled RNA-seq results depicted in Figures 5 and S5, as also deposited as ArrayExpress accession E-MTAB-7349.

## Supplementary Methods – D’Agati et al

### Vectors and transgenic lines

*The transgene vector col2a1aR2:KalTA4,myl7:EGFP* was assembled by Multisite Gateway Cloning combining *pENTR5’-col2a1aR2, pENTR’D-KalTA4 (1*), *p3E_SV40polyA (*Tol2kit *#302*) (*2*), and *pDestTol2CG2 (*Tol2kit *#395*) (*2*). Successful recombination was confirmed by restriction digest and Sanger sequencing. Wildtype embryos of the *TÜ* strain were co-injected with 25 pg of the final plasmid and 25 pg of *Tol2 transposase* mRNA at the one cell stage and raised according to standard procedures (*3*). F2 animals with a single-copy transgene insertion were selected for experimentation. The lines *Tg(-2.7twhh:Gal4)* and *Tg(5xUAS:eGFP-HRASV12) were previously described (4, 5*).

*UAS* vectors for overexpression of candidate genes were also generated by Multisite Gateway Cloning. The ORF of chordoma candidate genes was amplified from human cDNA clones or zebrafish cDNA (see below) using the primers listed in Table 1 and TOPO-cloned into *pENTR/D-TOPO (*Thermo Scientific) according to the manufacturer’s instructions. Zebrafish *kdr/vegfr2* follows suggested ortholog nomenclature (*6, 7*). Gateway reactions were performed with *pENTR5’-4xnr UAS (8*), *pAF20-3’-2AmCerulean (9*) and *pCM326 (pDestTol2CG2,crya:Venus*) (*10*). *Tg(col2a1aR2:KalTA4,myl7:EGFP)* embryos were injected with 25 pg of the *pUAS:candidate* together with 25 pg of *Tol2 transposase* mRNA at the one cell stage and raised until 5 dpf (*3*).

**Table 1:**
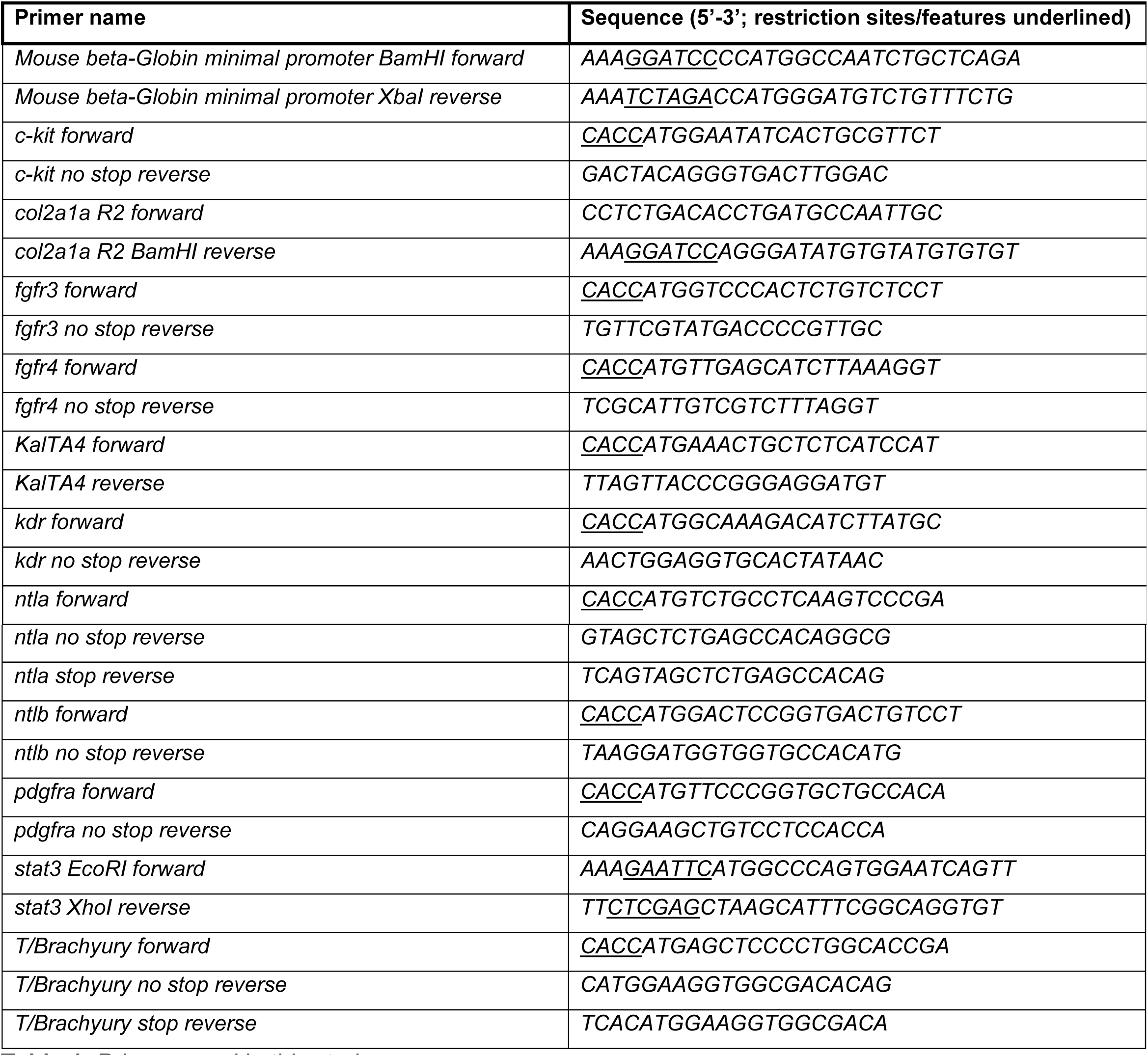
Primers used in this study.

### Primers and vectors

**Table 2:** List of Plasmids used in the transient injection approach

| Plasmid name | Overexpressed ORF |
| --- | --- |
| *UAS:EGFR-2AEGFP* | *EGFR* (human), fusion protein with 2A-EGFP ORF |
| *UAS:fgfr3-2AEGFP* | *fgfr3* (zebrafish), fusion protein with 2A-EGFP ORF |
| *UAS:fgfr4-2AEGFP* | *fgfr4* (zebrafish), fusion protein with 2A-EGFP ORF |
| *UAS:EGFP-HRAS<sup>V12</sup>* | *HRAS<sup>G12V</sup>* (human), direct fusion with *EGFP* ORF |
| *UAS:kdr-2Acerulean (vegfr2)* | *kdr/vegfr2* (zebrafish), fusion protein with 2A-EGFP ORF |
| *UAS:c-kit-2AEGFP* | *kita/c-kit* (zebrafish), fusion protein with 2A-EGFP ORF |
| *UAS:ntla* | *ntla* (zebrafish) |
| *UAS:ntla-2AEGFP* | *ntla* (zebrafish), fusion protein with 2A-EGFP ORF |
| *UAS:ntla-VP16* | Zebrafish *ntla aa1-232+VP16* transactivation domain |
| *UAS:ntlb* | *ntlb* (zebrafish) |
| *UAS:ntlb-2AEGFP* | *ntlb* (zebrafish), fusion protein with 2A-EGFP ORF |
| *UAS:pdgfra-2AEGFP* | *pdgfra* (zebrafish), fusion protein with 2A-EGFP ORF |
| *UAS:EGFP-rheb* | *rheb* (zebrafish) |
| *UAS:EGFP-stat3* | *stat3* (zebrafish) |
| *UAS:Brachyury* | *Brachyury* (human) |
| *UAS:Brachyury-2AEGFP* | *Brachyury* (human), fusion protein with 2A-EGFP ORF |
| *UAS:EGFP* | *EGFP* |

### Antibodies

Antibodies used in this study comprise anti-pERK (4376, Cell Signaling, Danvers, MA; USA), anti-Cytokeratin (961, Abcam, Cambridge, UK), anti-TP53 (GTX128135, GeneTex, Irvine, CA, USA) and anti-Ntla (a gift from Andy Oates, EPFL) (*11*) and anti-Brachyury (sc-20109 or sc-166962; Santa Cruz, Dallas, TX).

RNA-Seq reads were mapped to the zebrafish reference genome (*GRCz10*) using STAR (*12*). Mapped reads were then quantified with featureCounts (*13*). We then used edgeR (*14*) for differential expression analysis with a FDR cutoff of 5%. Pathway analysis was done using Ingenuity Pathway Analysis software (https://www.qiagenbioinformatics.com/products/ingenuity-pathway-analysis/) and the Gene Set Enrichment Analysis packages (http://software.broadinstitute.org/gsea/index.jsp) using the log2FC and FDR < 0.05. The RNA-seq data is available through ArrayExpress accession E-MTAB-7349 and summarized as Excel table in Table S1.

